# Weak electric fields promote resonance in neuronal spiking activity: analytical results from two-compartment cell and network models

**DOI:** 10.1101/379560

**Authors:** Josef Ladenbauer, Klaus Obermayer

## Abstract

Transcranial brain stimulation and evidence of ephaptic coupling have sparked strong interests in understanding the effects of weak electric fields on the dynamics of neuronal populations. While their influence on the subthreshold membrane voltage can be biophysically well explained using spatially extended neuron models, mechanistic analyses of neuronal spiking and network activity have remained a methodological challenge. More generally, this challenge applies to phenomena for which single-compartment (point) neuron models are oversimplified. Here we employ a pyramidal neuron model that comprises two compartments, allowing to distinguish basal-somatic from apical dendritic inputs and accounting for an extracellular field in a biophysically minimalistic way. Using an analytical approach we fit its parameters to reproduce the response properties of a canonical, spatial model neuron and dissect the stochastic spiking dynamics of single cells and large networks. We show that oscillatory weak fields effectively mimic anti-correlated inputs at the soma and dendrite and strongly modulate neuronal spiking activity in a rather narrow frequency band. This effect carries over to coupled populations of pyramidal cells and inhibitory interneurons, boosting network-induced resonance in the beta and gamma frequency bands. Our work contributes a useful theoretical framework for mechanistic analyses of population dynamics going beyond point neuron models, and provides insights on modulation effects of extracellular fields due to the morphology of pyramidal cells.

**Author Summary:** The elongated spatial structure of pyramidal neurons, which possess large apical dendrites, plays an important role for the integration of synaptic inputs and mediates sensitivity to weak extracellular electric fields. Modeling studies at the population level greatly contribute to our mechanistic understanding but face a methodological challenge because morphologically detailed neuron models are too complex for use in noisy, in-vivo like conditions and large networks in particular. Here we present an analytical approach based on a two-compartment spiking neuron model that can distinguish synaptic inputs at the apical dendrite from those at the somatic region and accounts for an extracellular field in a biophysically minimalistic way. We devised efficient methods to approximate the responses of a spatially more detailed pyramidal neuron model, and to study the spiking dynamics of single neurons and sparsely coupled large networks in the presence of fluctuating inputs. Using these methods we focused on how responses are affected by oscillatory weak fields. Our results suggest that ephaptic coupling may play a mechanistic role for oscillations of population activity and indicate the potential to entrain networks by weak electric stimulation.

## Introduction

The interaction between weak electric fields and neuronal activity in the brain has gained increased attention over the past decade [1]. These weak fields can be generated endogenously by populations of neurons [2–4] or through transcranial electrical stimulation [5–7], and they can modify neural activity in various ways [2,8–11]. Although the electric fields caused by this type of noninvasive intervention exhibit low magnitudes (≤ 1-2 V/m [5, 6]) they can modulate neuronal spiking activity [2, 12, 13] and lead to changes in cognitive processing [14, 15], offering a number of possible clinical interventions [16, 17]. The influence of extracellular fields on the subthreshold membrane voltage of single cells has been thoroughly studied and biophysically explained [12, 18–21]. How weak electric fields affect neuronal spiking activity and interact with network dynamics, however, is currently not well understood.

Multi-compartment models are useful tools to dissect effects in single neurons in the absence of input fluctuations [22–24], but they are not well suited to study neuronal and network spiking activity in noisy, in-vivo like conditions, because of their large complexity. Single-compartment (point) neuron models on the other hand allow for effective mechanistic analyses at the network level (see, e.g., [25–27]); however, they lack the spatial structure that is required to biophysically describe the effects of an electric field [20]. Nevertheless, networks of point model neurons have been repeatedly used in conjunction with rough phenomenological implementations of extracellular field effects [2, 9, 11, 28].

Here we employ a model of pyramidal (PY) spiking neurons with the minimal level of spatial detail necessary to biophysically take into account an extracellular electric field. It consists of a compartment for the soma and one for the apical dendrite, allowing to differentiate between synaptic inputs at the soma (including basal dendrite) and those at the distal (apical) dendrite. The model parameters are semi-analytically tuned to reproduce the behavior of a more sophisticated, spatially extended neuron model which involves the cable equation. To effectively study neuronal spiking dynamics for in-vivo like fluctuating inputs and the activity of sparsely coupled large populations in the presence and absence of extracellular fields we develop an analytical method. It utilizes the Fokker-Planck equation and a moment closure approximation technique to characterize the stochastic model dynamics.

Using these tools we study (i) how a weak oscillatory field affects the spiking activity of neurons exposed to fluctuating synaptic inputs, (ii) how these effects compare to those of weak oscillatory inputs in the absence of an electric field, and (iii) how weak applied fields modulate network-induced oscillations. Our contribution sheds some light on the effects of extracellular fields at the population level. Furthermore, it provides useful methods for mechanistic studies on the dynamics of coupled compartmentalized spiking neurons that allow to broadly distinguish inputs according to the location of the synapses. Such a distinction is important, for example, in circuit models which involve different types of (inhibitory) neurons.

## Results

### Modeling approach

#### Pyramidal neuron model

The PY model neurons consist of two compartments, one for the soma and one for the (apical) dendrite, for which we consider trans-membrane capacitive currents, ionic leak currents, an approximation of the somatic Na^+^ current at spike initiation, an internal current, synaptic input currents and an extracellular field. The latter is defined as 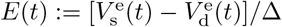, where 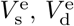 are the extracellular membrane potentials for the somatic and dendritic compartments, whose centers are separated with distance Δ. The dynamics of action potentials are simplified by a reset mechanism of the integrate-and-fire type at the soma. Fluctuating input currents at the soma and the dendrite, *I*_s_ and *I*_d_, that mimic the compound effect of synaptic bombardment in vivo, are described by stochastic processes with means *Ī*_s_, *Ī*_d_ and standard deviations *σ*_s_, *σ*_d_. In this model the dynamics of the somatic and the dendritic membrane voltage, *V*_s_ and *V*_d_, respectively, are thus governed by two coupled differential equations together with a reset condition for spikes (for details see Methods section 1). A schematic circuit diagram of the model is depicted in Fig 1A.

**Figure 1.**
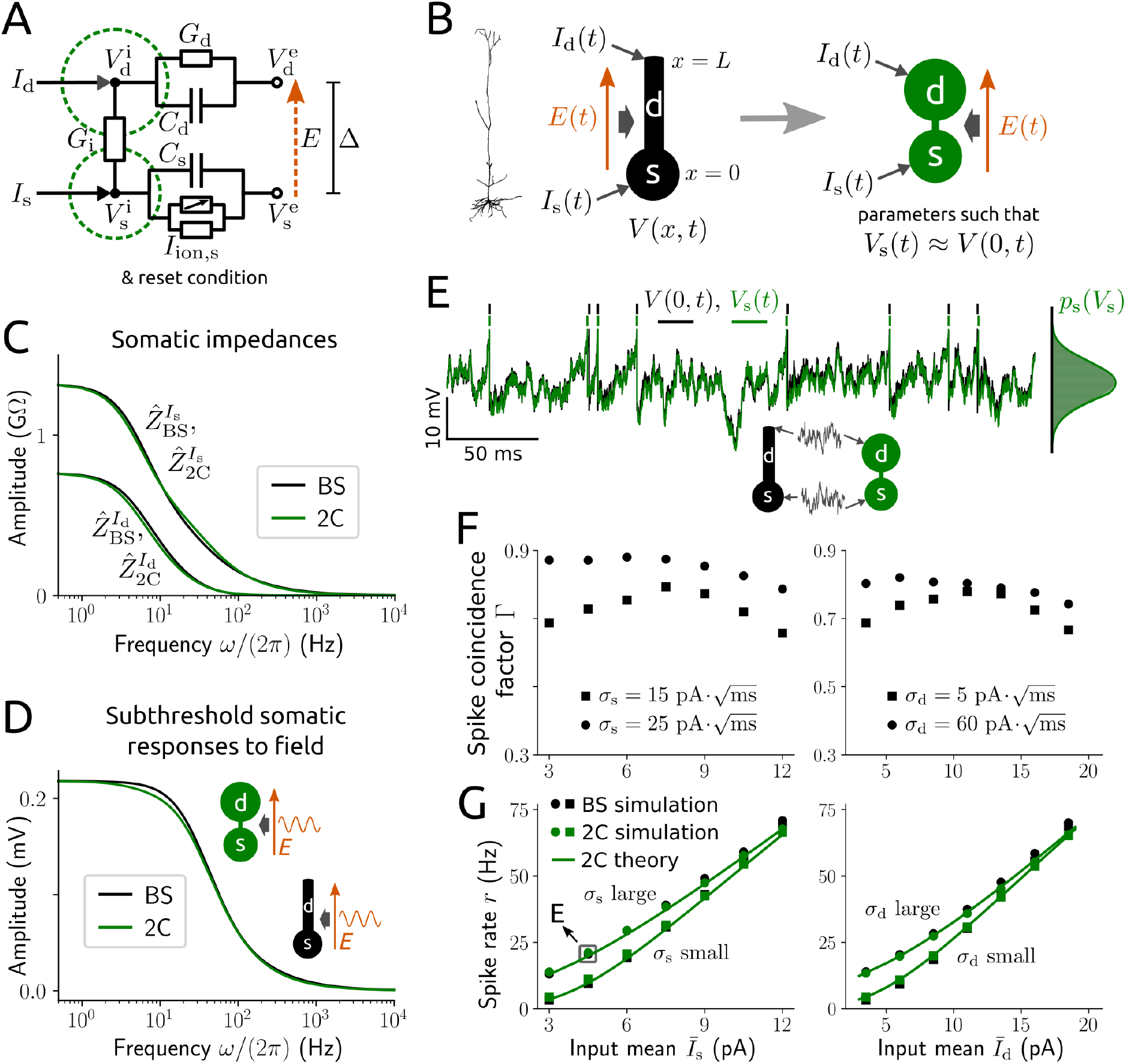
PY neuron model: parameter fitting and response properties. **A**: schematic circuit diagram for the membrane voltage dynamics of the two-compartment (2C) model (for the full description see Methods section 1). **B**: visualization of a spatial ball-and-stick (BS) model neuron (black) whose somatic voltage dynamics is approximated by the 2C model (green). *I*_s_, *I*_d_ denote the input currents at the soma and dendrite, *E* is the extracellular electric field and VS is the somatic membrane voltage of the 2C model neuron and *V* is the membrane voltage of the BS model. **C–G**: responses of a BS neuron parametrized to model a typical PY cell and of the fitted 2C model (for parameter values see Table 1). **C**: amplitude of subthreshold somatic impedances for inputs at the soma *Ẑ*^*I*_s_^ and dendrite *Ẑ*^*I*_d_^ as a function of input frequency (using Eqs (11), (12) and (67)–(69)). **D**: amplitude of subthreshold somatic voltage responses to a sinusoidal electric field with amplitude *E*_1_ =1 V/m (and *E*_0_ = 0) as a function of field frequency. **E**: example time series of the somatic voltage in response to fluctuating inputs, with spike times indicated, as well as voltage histograms of both models (right) and voltage density *p_s_* (green line; analytically calculated, cf. Methods section 3). **F**: spike coincidence measure Γ between the spike trains of both models for *Ī*_d_ = 3 pA, 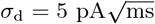 (left) and *Ī*_s_ = 3 pA, 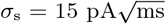 (right). Γ = 1 indicates an optimal match with precision Δ_c_ = 5 ms, Γ = 0 indicates pure chance (for details see Methods section 5). **G**: spike rates from numerical simulation (symbols) and analytically calculated (green line; cf. Methods section 3) for input parameter values as in **F**. The grey square marks the parameter values used for **E**.

#### Model parametrization

To determine the parameter values of the two-compartment model and assess whether its spatial complexity is adequate we semi-analytically fit the model to a biophysically more sophisticated, spatially elongated ball-and-stick neuron model (Fig 1B). That model involves the cable equation, an integrate-and-fire spike mechanism at the soma and an extracellular field which is assumed spatially homogeneous (see Methods section 4). Due to its mathematical complexity this model is not well suited for analyses of spiking dynamics and application in networks. The fitting method approximates the somatic responses of the ball-and-stick model in an efficient way without requiring the parameter values for the input or the electric field. In brief, we analytically calculate the somatic subthreshold responses for small amplitude variations of the inputs and the electric field as well as the transient somatic voltage responses after a spike for threshold inputs, and subsequently apply a least-squares fit. Note that the membrane voltage dynamics at the soma directly affect spike timing and are therefore the most relevant (axon initial segments are absorbed by the somatic compartments and not separately included in the models). In this way we rapidly obtain an accurate reproduction of the relevant response properties of the ball-and-stick model (Fig 1C-G). Specifically, the subthreshold responses, in terms of somatic impedances for inputs at the soma and dendrite (Fig 1C), the somatic voltage response for an oscillatory applied field (Fig 1D), and spiking activity, in terms of spike coincidences (Fig 1F, see Methods section 5) and the spike rate (Fig 1G), are well approximated.

#### Characterization of spiking dynamics

We focus on spiking activity, in particular the dynamics of the (instantaneous) spike rate *r*. This quantity can be calculated exactly using the Fokker-Planck equation that governs the evolution of the joint probability density for the somatic and dendritic membrane voltage *p*(*V*_s_, *V*_d_,*t*), describing the stochastic dynamics in deterministic form (see Methods section 3). Since this equation cannot be solved numerically in reasonable time we employ a moment closure approximation method. Specifically, we use *p*(*V*_s_, *V*_d_,*t*) = *p*_s_(*V*_s_,*t*)*p*_d_(*V*_d_|*V*_s_,*t*), where *p*_s_ is the marginal probability density for the somatic voltage and *p*_d_ the probability density for the dendritic voltage conditioned on *V*_s_, and approximate *p*_d_ by a conditioned Gaussian probability density. The resulting system allows for convenient and efficient calculation of the spike rate responses for constant input statistics as well as weak sinusoidal variations of the input moments or the applied field. An evaluation of this method in comparison to numerical simulation for constant input moments is shown in Fig 1E,G.

### Modulation of neuronal spiking activity

We first consider a PY neuron exposed to fluctuating inputs at the soma and the dendrite and a weak sinusoidal applied field. The noisy inputs drive the neuron to stochastic spiking activity that is influenced by the field. This effect can be quantified by the instantaneous spike rate across a large number of trials (obtained from numerical simulations with different realizations of the input), which is equivalent to the population-averaged spike rate for a large number of uncoupled PY neurons receiving independent inputs. The field leads to an oscillatory modulation of the spike rate that is accurately reproduced by our analytical calculation method (Fig 2A).

**Figure 2.**
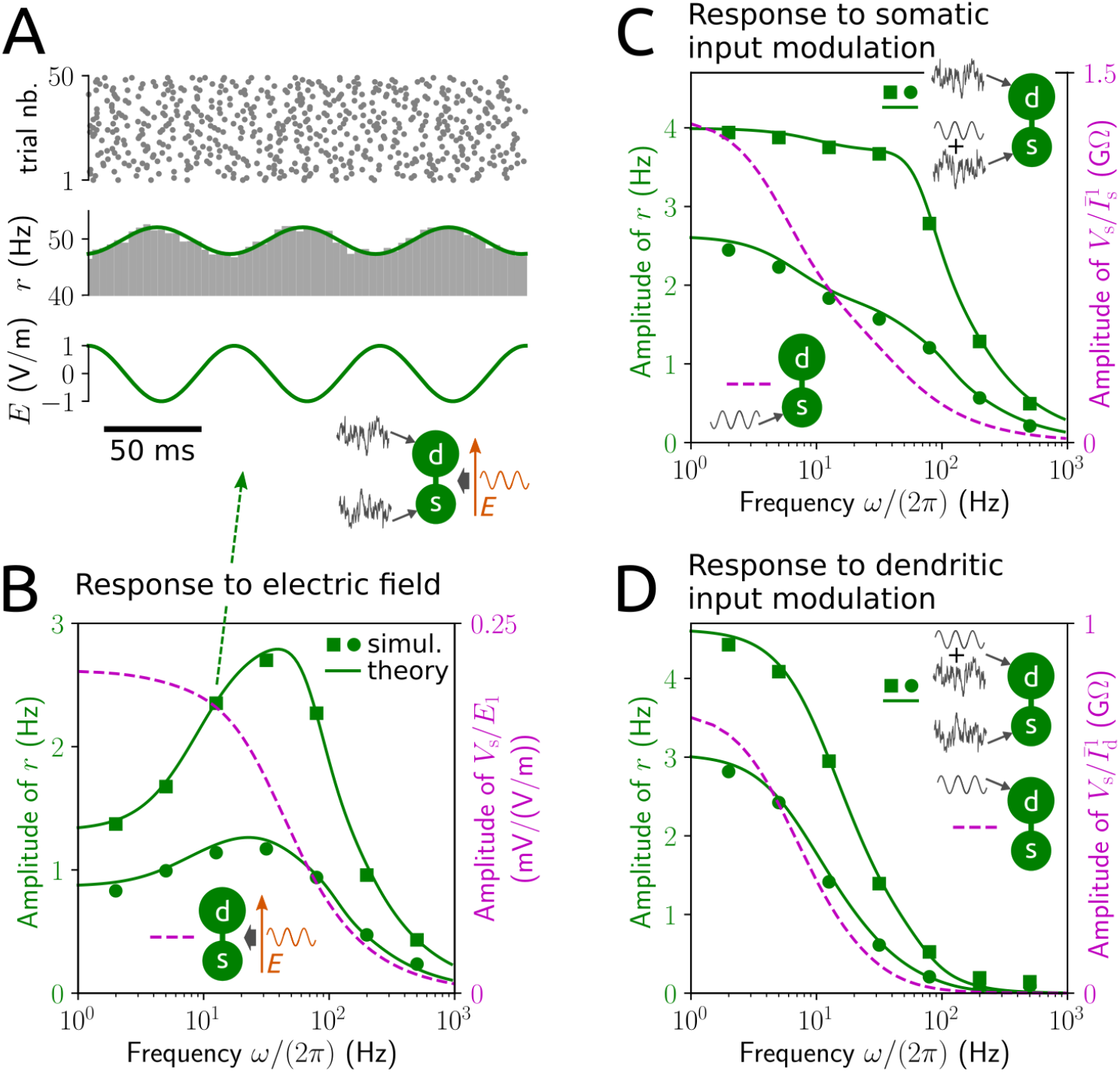
Neuronal responses to an applied electric field and to input modulations. **A**: spike times and spike rate (histogram: grey, analytical result: green line; cf. Methods section 3) of a PY model neuron in response to a sinusoidal electric field, *E*(*t*) = *E*_1_ sin(*ωt*) with angular frequency *ω* (here: *ω*/(2*π*) = 12.6 Hz), in the presence of noisy background input. **B**: amplitude of spike rate responses to a field with amplitude *E*_1_ = 1 V/m as a function of field frequency for *Ī*_s_ = 10 pA, 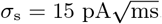, *I*_d_ =3 pA, 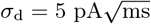 (green squares) and *Ī*_s_ =3 pA, 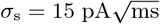, *Ī*_d_ = 7 pA, 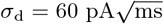 (green circles), as well as amplitude of normalized subthreshold somatic voltage responses (magenta dashed line). **C** and **D**: amplitude of spike rate responses (green) and of normalized subthreshold somatic voltage responses (impedances; magenta dashed lines) to sinusoidal modulations of the mean input at the soma, 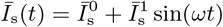, with amplitude 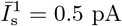 (**C**) or at the dendrite, 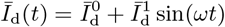, with amplitude 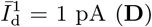 (**D**) as a function of modulation frequency in the absence of an extracellular field. Baseline (constant) input statistics in **C** and **D** correspond to those in **B**.

By measuring these spike rate responses over a range of field frequencies we observe a clear resonance in a biologically relevant, relatively narrow frequency band (Fig 2B). In other words, the spike rate oscillations are strongest for field oscillations in that frequency range. This resonance behavior is not shown by the subthreshold membrane voltage response to the field in the absence of suprathreshold fluctuating inputs. Interestingly, spike rate responses to weak sinusoidal modulations of the mean input at the soma or dendrite instead of an applied field do not exhibit such a resonance (Fig 2C,D). Response amplitudes typically decrease as the frequency of the input modulations increases, although spike rate responses remain elevated for somatic modulations of up to about 100 Hz. Notably, the analytical results are in good agreement with those of numerical simulations. This justifies the extensive application of our analytical method in the subsequent analyses.

Next, we assess the resonance behavior caused by the applied field for a range of biologically plausible input statistics (see Fig 3). Resonance frequency and strength increase with increasing input mean, but not necessarily with increasing input variance. For mean-dominated input (that is, large input mean and small input variance) the resonance frequency is similar to the baseline spike rate (compare the dashed curves for large input mean with Fig 2G). In this case an increase in input variance interestingly leads to a decrease in resonance frequency and strength. For fluctuation-dominated input the resonance frequency is restricted to the the beta and low gamma frequency bands.

**Figure 3.**
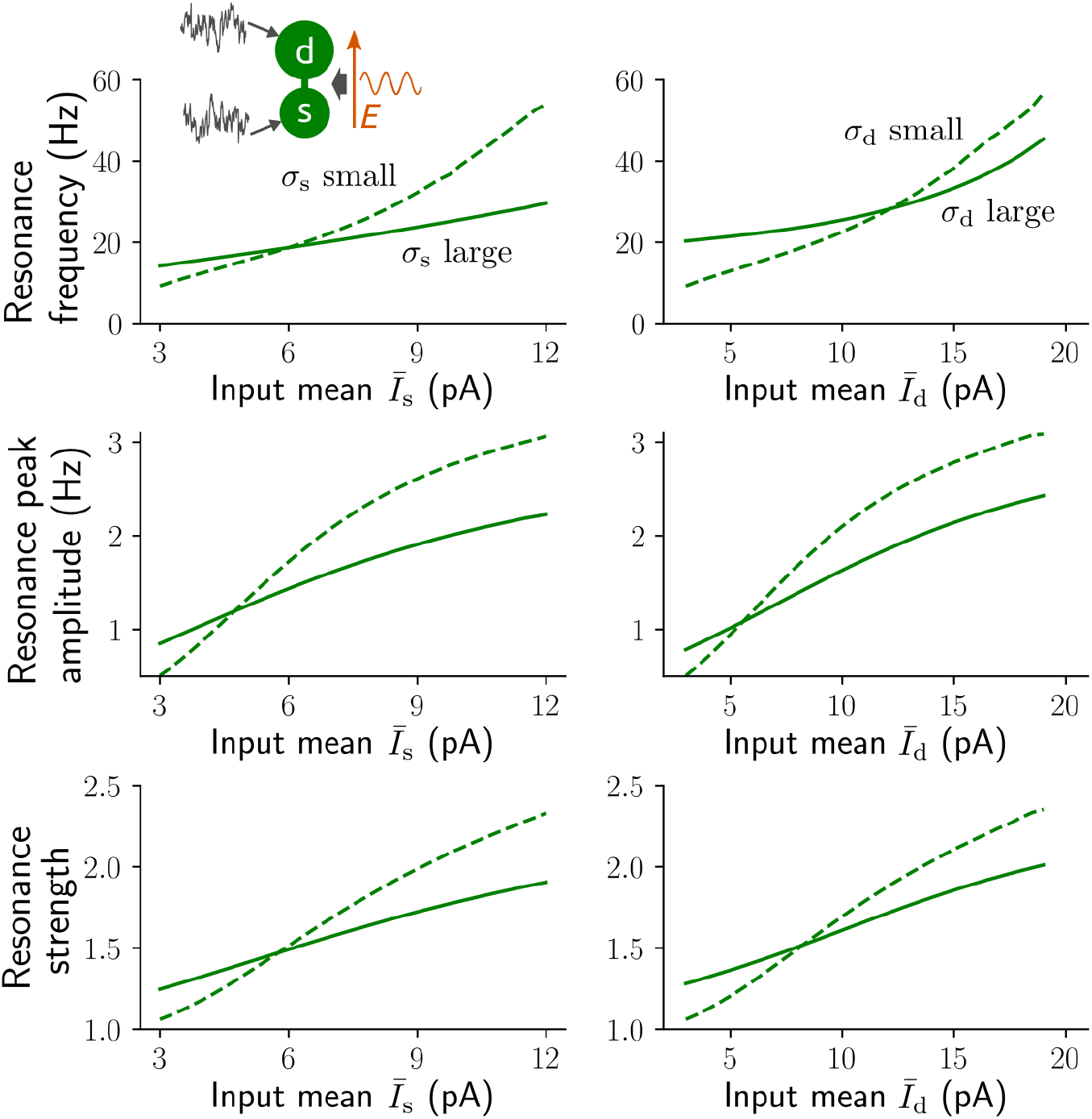
Spike rate resonance caused by an applied field. Resonance frequency (argmax_*ω*_ *r*_1_(*ω*)/(2*π*); top panel), amplitude (max*r*_1_(*ω*); center panel) and strength (max*r*_1_(*ω*)/*r*_1_(0); bottom panel) of the oscillatory spike rate with amplitude *r*_1_ (cf. Methods section 3) in response to a sinusoidal applied field with *E*_1_ = 1 V/m and angular frequency *ω* for a range of input statistics. Left: 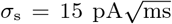 (dashed lines), 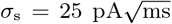 (solid lines), *Ī_d_* = 3 pA and 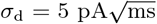 right: 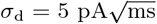 (dashed lines), 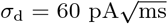 (solid lines), *Ī*_s_ = 3 pA and 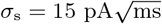.

How does the external field promote this resonance behavior? From the definition of the extracellular field and the circuit diagram in Fig 1A it becomes evident that the trans-membrane currents caused by the field at the soma and dendrite are opposed (using Kirchhoff’s law, that all incoming currents at a point of the circuit must sum to zero; see Eqs. (1), (2) in Methods section 1). This indicates that the electric field effectively reflects anti-correlated inputs at the soma and dendrite. To examine the influence of input correlations more closely we applied sinusoidal modulations of the mean input at the soma and dendrite with a phase shift (Fig 4A). A resonance peak emerges as the phase shift increases from 0 (synchronized modulations / strong correlation) towards 180° (anti-synchronized modulations / negative correlation) and becomes most pronounced at that value. To explore the effects of synaptic delays in this regard we further considered a temporal shift instead of a phase shift between the two mean input modulations (Fig 4B). Delays that are sufficiently large cause multiple resonance peaks. The frequency that corresponds to the most dominant peak (with positive frequency) decreases with increasing delay, but is otherwise largely independent of the baseline input statistics (Fig 4C).

**Figure 4.**
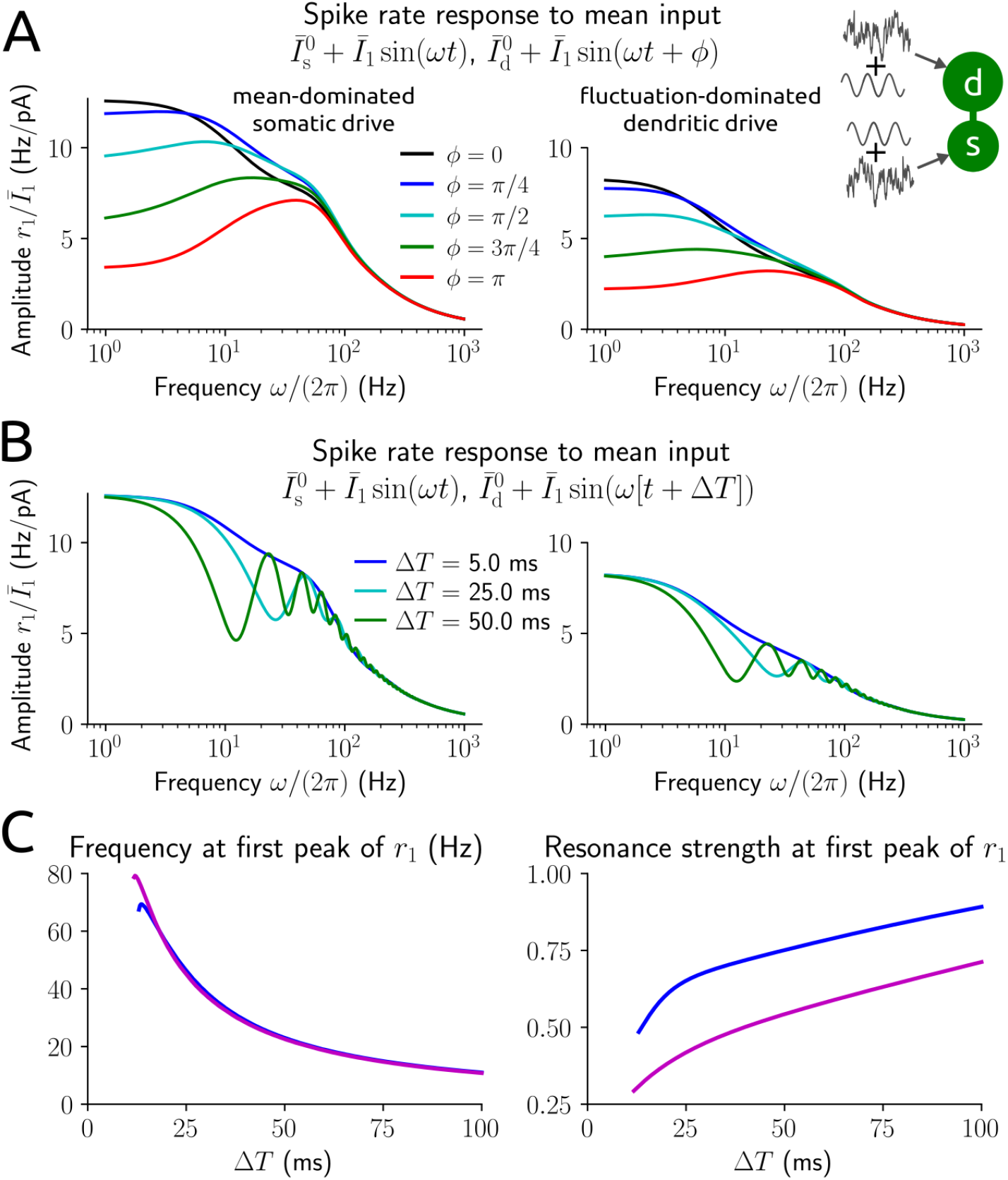
Spike rate responses to input modulations at soma and dendrite. **A**: amplitude of normalized rate responses to sinusoidal modulations of the mean input at the soma, 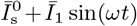, and dendrite, 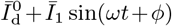, with phase shift *ϕ* as a function of modulation frequency for the baseline input statistics 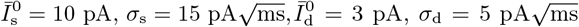 (left) and 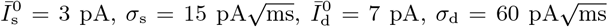 (right). *ϕ* = *π* corresponds to negatively correlated (anti-synchronous) modulations. **B**: amplitude of normalized rate responses to mean input modulations 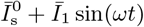 and 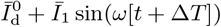 with temporal shift (delay) Δ*T*. Baseline input statistics as in **A. C**: frequency (left) and normalized amplitude (resonance strength, using the amplitude at frequency 0; right) of the most dominant peak at a positive frequency >1 Hz as a function of delay Δ*T* for baseline input statistics as in **A** (left: blue lines, right: magenta lines).

### Modulation of network dynamics

To examine how an applied field interacts with network mechanisms to shape population dynamics we derived a mean-field network model from large populations of sparsely coupled PY neurons and inhibitory (IN) interneurons exposed to fluctuating background inputs and a spatially homogeneous (with respect to neuronal orientation) weak electric field. PY neurons are described by the two-compartment model and IN neurons by an established single-compartment spiking model (exponential integrate-and-fire), because IN neurons are spatially more compact and therefore the direct effect of the electric field on them is negligible. Synaptic coupling is incorporated by delayed current pulses which cause post-synaptic potentials of reasonably small magnitude with distributed delays. In the derivation we apply a diffusion approximation and a moment closure method to transform the original network model into a manageable system of two Fokker-Planck equations – one for each population – coupled via synaptic input moments. This system describes the collective stochastic dynamics in a way that allows for convenient dissection of the network activity in terms of population-averaged instantaneous spike rates (see Methods section 6).

The network is parametrized to exhibit a baseline state of low or high asynchronous (irregular) spiking activity depending on the strength of the external drive. Network-induced resonance emerges for weak sinusoidal input modulations at the soma of PY neurons with strongest responses in the gamma frequency range (Fig 5). This resonance behavior is more pronounced in the high activity state compared to the low activity state. The resonance peak shifts to a higher or lower gamma frequency depending on whether IN neurons target PY neurons only at the soma or only at the dendrite. Interestingly, for input modulations at the dendrite of PY neurons such a clear resonance behavior does not appear.

**Figure 5.**
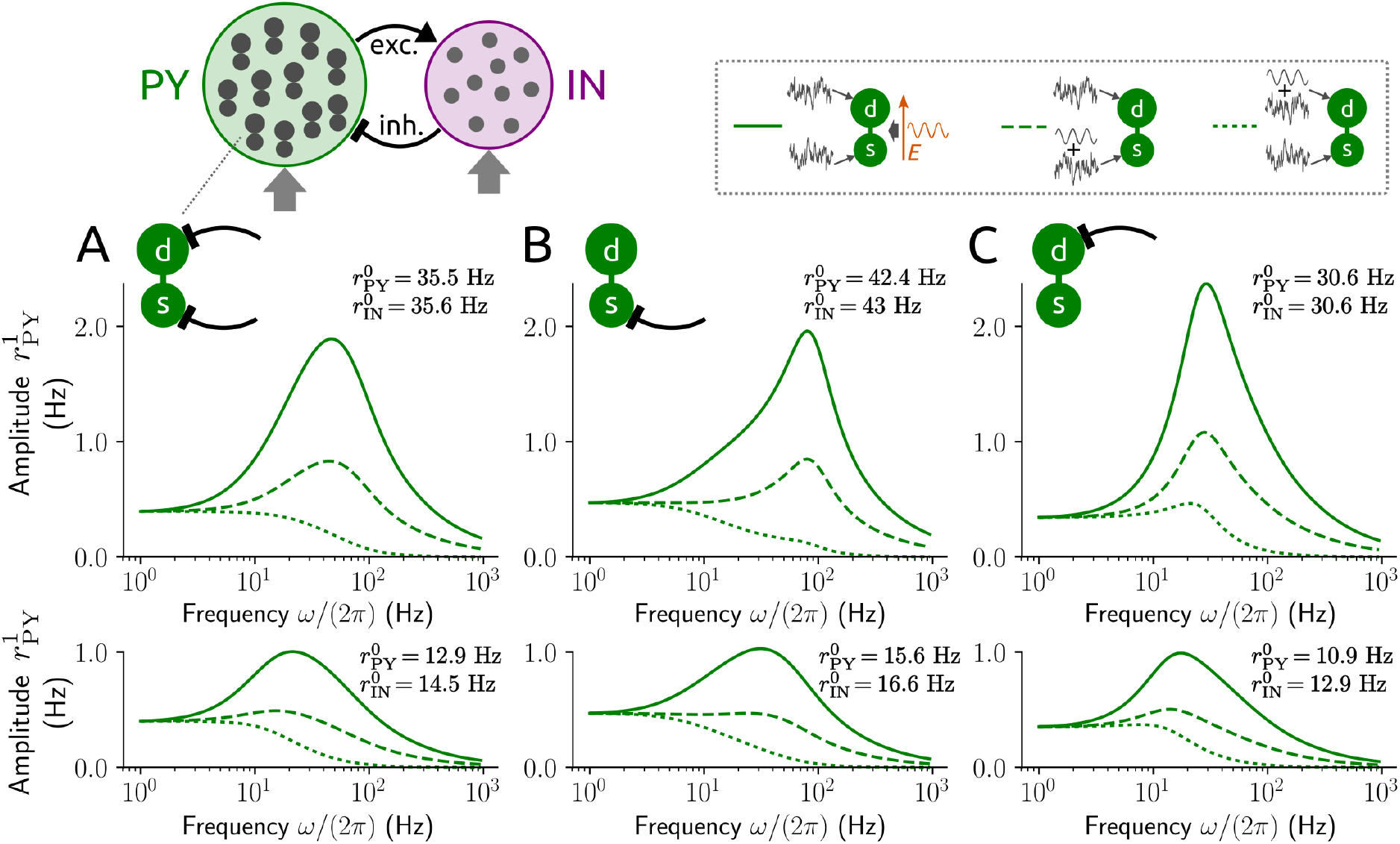
Resonance in PY-IN networks. Amplitude of oscillatory spike rate of the PY neuron population for a mean-field network of sparsely coupled PY and IN neurons exposed to an applied field (solid lines) or weak modulations of the mean input to PY neuron somas (dashed lines) and dendrites (dotted lines) as a function of field/modulation frequency. for strong (top panel) and weak (bottom panel) external drive. Each PY neuron receives inputs from 100 IN neurons both at soma and dendrite (**A**) or from 200 IN neurons only at the soma (**B**) or dendrite (**C**), each IN neuron receives inputs from 200 PY neurons; random connectivity, excitatory/inhibitory coupling strengths ±0.1 mV and distributed delays with bi-exponential delay density (rise time constant 0.5 ms, decay time constant 2 ms/5 ms for excitatory/inhibitory connections; for details see Methods section 6). Baseline mean of the external input for PY neurons (at soma and dendrite): 12 pA (top panel), 6 pA (bottom panel) and 0 pA for IN neurons; input standard deviation: 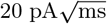 for PY neurons, 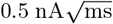 for IN neurons. Field amplitude was *E*_1_ = 1 V/m, input modulation amplitudes were chosen to yield equal response amplitude at the lowest frequency.

When we consider a weak sinusoidal field instead of input modulations we observe a strongly amplified resonance of the population activity in the same (gamma) frequency band. The prominent effect of an applied field in single cells thus carries over to PY-IN networks, boosting network-based oscillations (that are mediated by an excitation-inhibition loop). These results indicate that an oscillatory electric field as generated, for example, by transcranial stimulation should be most effective to induce or maintain rhythmic network dynamics in the gamma frequency range.

## Discussion

### Methodological aspects

#### Neuron model

Model-based investigations on how extracellular electric fields impact neuronal spiking activity face a methodological challenge: the neuronal spatial extent plays an important biophysical role, however, morphologically detailed neuron models are too complex for use in noisy, in-vivo like conditions, especially in large networks. Two-compartment neuron models provide a useful compromise between biophysical detail and analytical tractability in this regard. Models of this type have been applied to study, for example, the influence of dendritic morphology on neuronal responses in the absence of extracellular fields [29] and the effects of constant electric fields on the activity of single neurons and synchronization of neuronal pairs in the absence of input fluctuations [30, 31]. Our approach is based on a two-compartment spiking PY neuron model which accounts for an external electric field and includes fluctuating synaptic input at the soma and/or (apical) dendrite.

Using semi-analytical techniques this model was rapidly calibrated to well approximate the (subthreshold somatic and spiking) response properties of a morphologically more plausible ball-and-stick model neuron. Based on that spatially elongated model we have recently developed an extension for single-compartment (point) neuron models to match the subthreshold somatic responses to synaptic inputs and an applied weak electric field [20]. The methodology presented here entails several advantages compared to our previous work: the fitted neuron model involves a low-dimensional differential equation in contrast to an integro-differential equation (in case of the extended point model to account for dendritic filtering effects), allowing for faster numerical simulation, straightforward implementation of networks, and a natural way to account for an extracellular field. Notably, the latter feature highlights the main benefit compared to networks of point model neurons for which simple phenomenological input currents were used to implement field effects [2, 9, 11, 28]. Importantly, the two-compartment model further allowed for the application of analytical methods to study spiking dynamics.

#### Spike rate dynamics

We devised a method to efficiently study the spike rate dynamics of these two-compartment neurons and the population activity of sparsely coupled large networks. It employs the Fokker-Planck partial differential equation and a moment closure technique for dimension reduction which allows to numerically solve the resulting system in reasonable time. These solutions yield an accurate and efficient approximation of the instantaneous (population) spike rate (cf. Fig. 1G and Fig. 2).

Similar moment closure methods were previously applied in different contexts, such as integrate-and-fire point model neurons with synaptic dynamics (see, e.g., [32, 33] and references therein). In [32] the issue was raised that moment closure may not be applicable for certain ranges of parameter values. For the setting and regions of parameter space considered throughout the present study, however, this approximation method was well suited.

We approximated the conditional probability density *p*_d_(*V*_d_|*V*_s_,*t*) by a conditioned Gaussian, thereby closing the system of lower dimensional equations at the 3^rd^ central moment of *V*_d_ (assuming the 3^rd^ and higher cumulants are zero). Closure at lower order moments leads so markedly less accurate results due to similar timescales for the somatic and dendritic voltage and strong coupling between those variables. Closure at higher order moments, on the other hand, does not yield a substantial improvement but increases computational demands.

Notably, moment closure at order 1 is equivalent to the adiabatic approximation frequently applied for adaptive integrate-and-fire neurons in the presence of noise [27, 34–36]. In that case the conditional first centered moment (of the adaptation variable given membrane voltage) is approximated by the corresponding unconditioned moment. This usually leads to an accurate reproduction of the spike rate dynamics due to the circumstance that the timescales of the two variables are separated. Such an adiabatic approximation has also been applied for two-compartment Purkinje model neurons [29] which possess large dendritic trees. A substantial difference in somatic and overall dendritic capacitance justifies the assumption of separated timescales for those models. Our approach, in contrast, is also valid for model parametrizations that yield rather similar timescales for soma and dendrite, and therefore suitable for pyramidal cells.

It is worth noting that our analytical techniques also allow for correlated somatic and dendritic inputs (cf. Methods section 3); an investigation into input correlations, however, is beyond the scope of the present study. The methodological results presented here may further be used to derive a simple spike rate model in form of a low-dimensional differential equation from two-compartment model neurons. This could be achieved by adapting available approximation methods [27] to our reduced description based on the Fokker-Planck equation.

### Described phenomena

By applying our analytical tools we showed that oscillatory weak electric fields strongly modulate neuronal spiking in a rather narrow frequency band. The resonance frequency depends on the input statistics. Spike rate modulation exhibited a clear resonance in the beta and gamma frequency bands in particular for strongly fluctuating inputs.

Recent modeling results suggest that this resonance frequency may be largely determined by the location of background input (soma vs. distal dendrite) [20]. Our results confirm that resonance frequencies are higher for mainly dendritic background inputs compared to mainly somatic background inputs (cf. Fig. 3, top panel). However, unless the input at one location is completely extinguished (as in [20]) the statistics of background inputs at either location appear to play the dominant role. It may also be noted at this point that spike rate resonance frequencies are lower for model neurons that include an exponential nonlinearity (at the soma) compared to purely leaky integrate-and-fire neurons [20].

To date, experimental evidence for frequency-dependent modulation of neuronal activity by extracellular fields is very sparse (see [37] for a review). Weak electric fields alternating at 30 Hz have been shown to increase spiking coherence of pyramidal cells in rat hippocampal slices [13], and fields with high-frequency components have been evidenced to entrain spiking activity in ferret primary visual cortex more effectively than fields that only contain low-frequency components [2]. To the best of our knowledge, the effects of multiple field frequencies have not yet been experimentally assessed. Therefore, our results on spike rate resonance are currently not completely confirmed and may be regarded as predictions.

Interestingly, these resonance effects at the suprathreshold level are not shown by the subthreshold membrane voltage, whose amplitude monotonically decreases with increasing field frequency (cf. Fig. 1D, Fig. 2B). The latter phenomenon is in accordance with electrophysiological observations: the subthreshold response amplitude (which scales linearly with the field amplitude [18]) is of the same order of magnitude as that measured in pyramidal cells and decreases with increasing field frequency [12]. Note that the parameter values of our model were not optimized to reproduce this behavior quantitatively but can be adjusted accordingly in a straightforward way [20].

The two-compartment model naturally accounts for input filtering caused by the dendrite (for example, as described in [20]). Notably, the somatic impedance and spike rate response for input modulations at the soma exhibit rather distinct dependencies on input frequency (compare Fig. 1C and Fig. 2C, right), which may be attributed to the nonlinearity caused by spiking. The presence of the dendrite leads to increased spike rate responses to weak oscillatory input modulations with high frequency at the soma. This is consistent with previous findings for Purkinje neurons in vitro and using a two-compartment model similar to the one used here [29] despite marked differences in the morphology between Purkinje and pyramidal cells. Specifically, the ratio between dendritic and somatic membrane surface (hence, capacitance) are quite distinct for these two types of neurons, which explains certain differences in the results.

## Models and Methods

### 1. Two-compartment neuron model

The two-compartment neuron model consists of two differential equations for the dynamics of the somatic and the dendritic membrane voltage, *V*_s_ and *V*_d_, respectively, together with a reset condition of the integrate-and-fire type (for similar models with and without an extracellular field see [29, 31, 38, 39]),

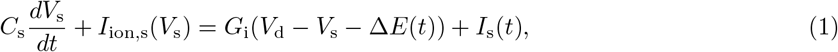

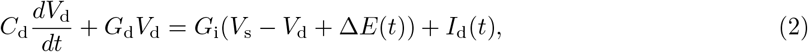

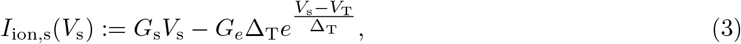

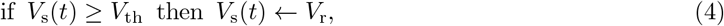

where *V*_s_, *V*_d_ are defined by the difference between the deviations 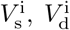 of the intracellular membrane potentials from the leak reversal potential (assumed identical for soma and dendrite) and the extracellular membrane potentials 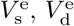 for the somatic and dendritic compartment, respectively,

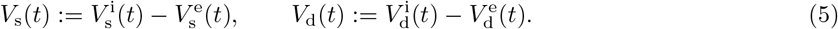

*C*_s_, *C*_d_ and *G*_s_, *G*_d_ denote the capacitances and leak conductances of the somatic and dendritic membranes. The exponential term with conductance *G_e_*, threshold slope factor Δ_T_ and effective threshold voltage *V*_T_ approximates the rapidly increasing Na^+^ current at spike initiation [40]. *G*_i_ is the internal conductance between the somatic and the dendritic compartment, Δ is the spatial distance between their centers, and *E* denotes the extracellular electric field, defined by

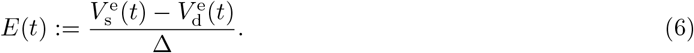

*I*_s_ and *I*_d_ are the synaptic input currents at the soma and dendrite, respectively. When *V*_s_ increases beyond *V*_T_, it diverges to infinity in finite time due to the exponential term, which defines a spike. In practice, however, the spike is said to occur when *V*_s_ reaches a given threshold value *V*_th_ > *V*_T_. The downswing of the spike is not explicitly modeled; instead, when *V*_s_ passes *V*_th_ (from below), the somatic membrane voltage is instantaneously reset to a lower value *V*_r_, cf. (4).

Eqs. (1) and (2) are the current balance equations for the center points of the two compartments according to the electrical circuit diagram in Fig. 1A. This can be seen by using Kirchhoff’s law, that all incoming currents at a circuit point must sum to zero, and the definitions (5) and (6) which imply 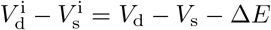. We consider an applied weak sinusoidal field,

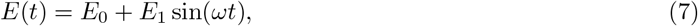

with offset *E*_0_ = 0, amplitude *E*_1_ = 1 V/m and angular frequency *ω*, unless stated otherwise.

The synaptic inputs are fluctuating currents that mimic the compound effect of synaptic bombardment in-vivo, described by

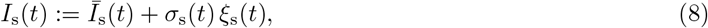

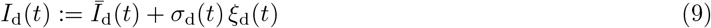

with time-varying moments *Ī*_s_, *Ī*_d_ and *σ*_s_, *σ*_d_, and uncorrelated unit Gaussian white noise processes *ξ*_s_, *ξ*_d_, i.e., 〈*ξ*(_s_(*t*)*ξ*_d_(*t* + *τ*)〉 = *δ*(*τ*)*δ*_sd_, where 〈·〈 denotes expectation (with respect to the ensemble of noise realizations at times *t* and *t* + *τ*) and *δ*_sd_ is the Kronecker delta.

### 2. Calculation of subthreshold responses

We analytically calculate the somatic membrane voltage response for small amplitude variations of the synaptic inputs *I*_s_(*t*), *I*_d_(*t*) and a weak oscillatory field *E*(*t*), which do not elicit spikes. Considering that the somatic voltage evolves sufficiently below the effective threshold value *V*_T_ allows us to neglect the exponential term in Eq. (1) (i.e., the somatic membrane is purely leaky and capacitive). Using the Fourier transform of Eqs. (1) and (2) we then obtain:

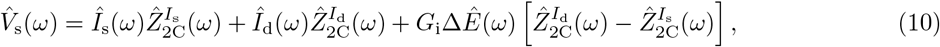

where

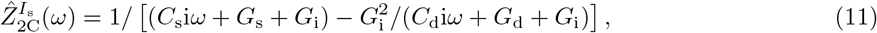

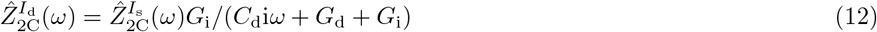

are the somatic impedances for inputs at the soma and at the dendrite, respectively. 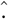 indicates the a Fourier transformed variable and *ω* denotes angular frequency.

### 3. Calculation of spike rate responses

To assess spiking activity we solved the stochastic differential equations Eqs. (1)–(4), (7)–(9) numerically using the Euler-Maruyama integration scheme with time steps between 0.01 ms and 0.05 ms. In addition, we employed analytical calculations as described in the following.

#### Fokker-Planck system

For improved readability we rewrite Eqs. (1)–(3) with extracellular field according to Eq. (7) and synaptic input given by Eqs. (8) and (9) in compact form:

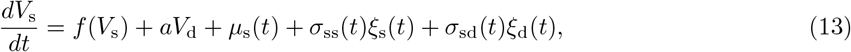

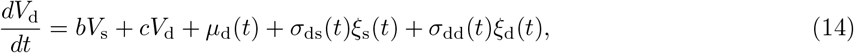

where the coefficients on the right hand side depend on the parameters of the system described in Methods section 1. Note that here *σ*_sd_ = *σ*_ds_ = 0, since the input fluctuations at the soma and dendrite are uncorrelated; however, the methods in this section may also be applied in scenarios where any of these parameters is nonzero and varies over time.

The dynamics of the joint membrane voltage probability density *p*(*V*_s_, *V*_d_,*t*) for this system plus reset condition (4) are governed by the Fokker-Planck equation (see, e.g., [27, 34, 41])

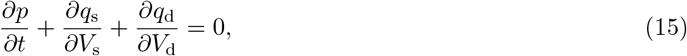

where *q*_s_ and *q*_d_ are the probability fluxes for the somatic and dendritic membrane voltage, respectively, given by

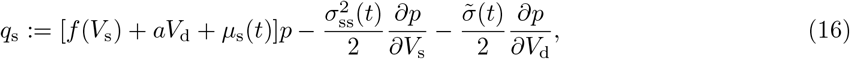

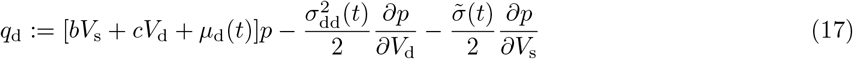

with 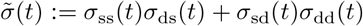, subject to the boundary conditions:

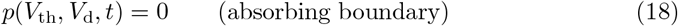

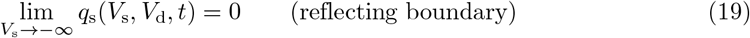

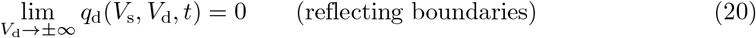

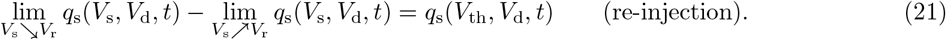

The latter condition, a re-injection of probability flux, accounts for the voltage reset in the neuron model. We obtain the instantaneous spike rate as

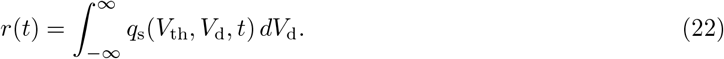

#### Dimension reduction

Solving the 2+1 dimensional Fokker-Planck partial differential equation (PDE) system (15)–(21) numerically is possible in principle, but computationally demanding. To reduce the dimension of this PDE system we utilize a moment closure approximation method. The (full) probability density *p* can be expressed in terms of the marginal probability density for the somatic voltage, *p*_s_, and the conditional probability density for the dendritic voltage, *p*_d_, as *p*(*V*_s_, *V*_d_, *t*) = *p*_s_(*V*_s_, *t*) *p*_d_(*V*_d_|*V*_s_, *t*). Note that *p*_d_ is characterized by a (potentially) infinite number of conditioned moments {*η*_d,1_ (*V*_s_, *t*), *η*_d,2_(*V*_s_, *t*),… }. The method approximates *p*_d_ by considering only the first *k* moments as described below (see [32] for the application of such a method in a different setting). We transform the PDE system (15)–(21) into a system of 1+1-dimensional PDEs

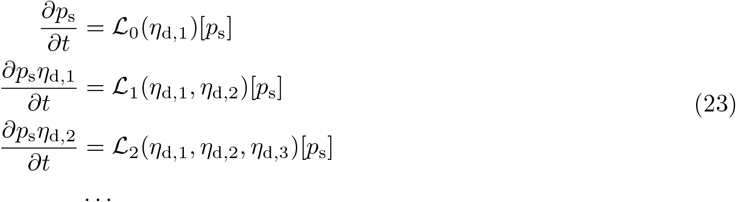

together with the associated boundary conditions by multiplying Eqs. (15)–(21) with 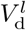 for *l* ∈ {0,1, 2,… } and integrating over *V*_d_ assuming that *p* and *q*_d_ tend sufficiently fast to zero for *V*_d_ → ±∞, i.e., 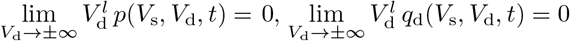. Each linear operator 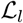 in (23) depends on the next higher conditioned moment *η*_d,*l*_+1, hence the system is (potentially) infinitely large. Note that we have omitted the obvious arguments *V*_s_,*t* for *p*_s_, *η*_d,*l*_ for improved readability.

We close the system (23) at *k* = 3 by setting the 3^rd^ central moment of *V*_d_ (as well as higher cumulants) to zero, such that 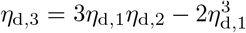, thereby assuming that *p*_d_ can be sufficiently well approximated by a conditioned Gaussian probability density,

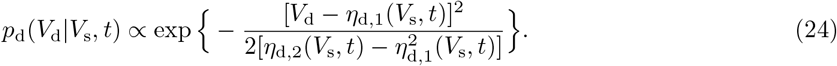

For a motivation of this assumption see the remark below. Defining *p*_s,1_:= *p*_s_*η*_d,1_ and *p*_s,2_:= *p*_s_*η*_d,2_ we obtain the system of 3 coupled PDEs:

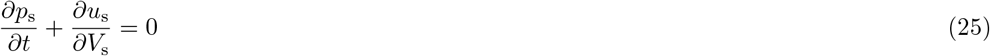

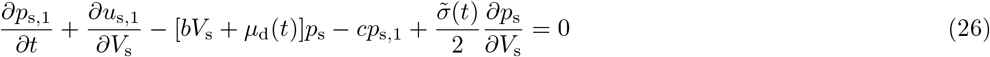

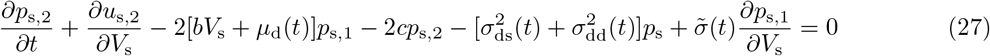

with

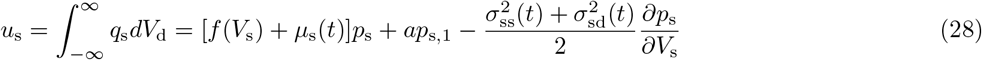

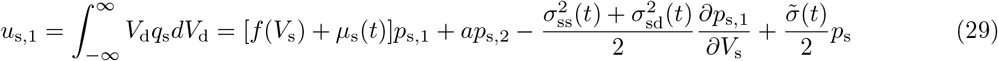

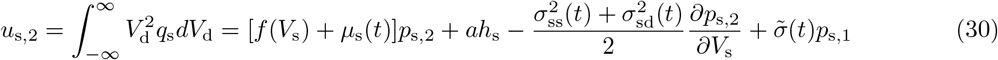

and

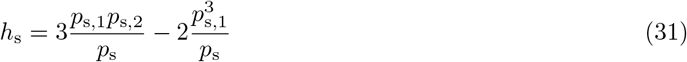

subject to the conditions

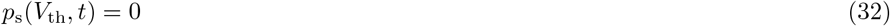

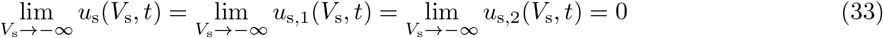

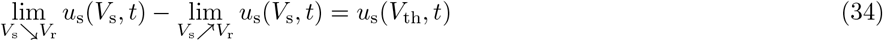

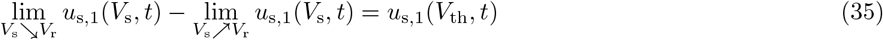

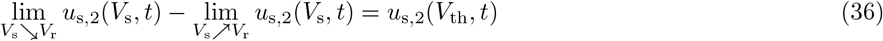

and requiring that *p*_s_ is initialized such that 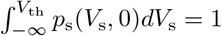. The spike rate is then obtained by

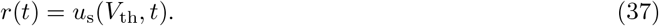

Note that *p*_s_(*V*_th_,*t*) = 0 implies *p*_s,1_(*V*_th_,*t*) = *p*_s,2_(*V*_th_,*t*) = 0. The conditions (33) follow from condition (19) and a self-consistency requirement with respect to the dynamics of the unconditioned moments 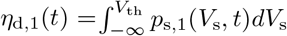 and 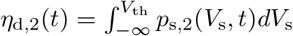. The latter can be seen by integration of Eqs. (26), (27) over *V*_s_ and comparison with the moment equations obtained by successive integration of Eq. (15) over *V*_s_, multiplication by *V*_d_ or 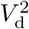, respectively, and integration over *V*_d_. Conditions (34)–(36) follow from (21). Note also that

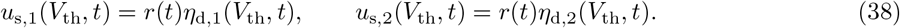

*Remark*: The assumption that *p*_d_ can be sufficiently well approximated by a conditioned Gaussian is supported by the circumstance that for subthreshold inputs and an electric field which keep the somatic voltage (sufficiently) below the spike threshold the approximation is excellent. In that case *p*_d_(*V*_d_|*V*_s_,*t*) is indeed a conditioned Gaussian probability density (because the exponential term in Eq. (1) as well as conditions (18) and (21) are negligible). Since this is not the case for stronger inputs (that cause spiking) the reproduction performance of this approximation needs to be evaluated (cf. Fig 1E,G, Fig 2 and Discussion).

#### Steady state

For the steady state (in case of constant parameters *μ*_s_,*μ*_d_, *σ*_ss_, *σ*_ds_, *σ*_sd_, *σ*_dd_) we obtain, by setting the time derivatives in Eqs. (25)–(27) to zero, the 6-dimensional ordinary differential equation (ODE) system

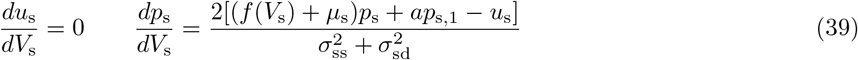

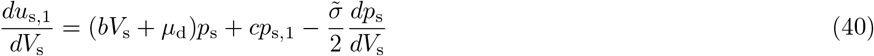

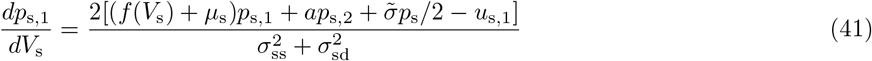

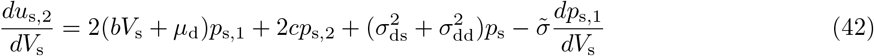

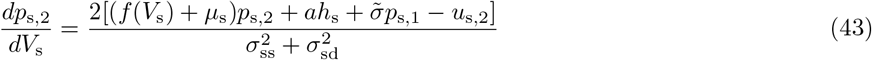

subject to the conditions (32)–(36) (with time dependence omitted).

*Numerical solution* We solve this nonlinear ODE system (nonlinearity due to *h*_s_, cf. Eq. (31)) with variable coefficients numerically by integrating Eqs (39)–(43) backwards from *V*_th_ with *p*_s_(*V*_th_) = *p*_s,1_(*V*_th_) = *p*_s,2_(*V*_th_) = 0, *u*_s_(*V*_th_) = 1, *u*_s,1_ (*V*_th_) = *η*_d,1_ (*V*_th_), *u*_s,2_(*V*_th_) = *η*_d,2_(*V*_t_h__) to a sufficiently small (lower bound) voltage value *V*_1b_, taking into account the “jump” conditions (34)–(36), and determine *η*_d,1_(*V*_th_) and *η*_d,2_(*V*_th_) such that *u*_s,1_(*V*_1b_) = *u*_s,2_(*V*_1b_) = 0, cf. condition (33), is fulfilled. Then, the scaling factor r (the spike rate) of the obtained solution is determined such that 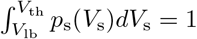 holds. The solution was achieved by means of Python implementation using a root finding algorithm provided by the package Scipy [42] (`optimize.root`, modification of the Powell hybrid method [43]) and low-level virtual machine acceleration through the package Numba [44].

#### Response to modulations

In order to characterize the spike rate dynamics we calculate the response to small-amplitude sinusoidal variations of the mean input around a baseline, 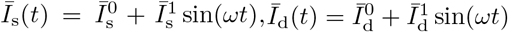, or to a weak sinusoidal field, cf. Eq. (7). These modulations translate to sinusoidal modulations of *μ*_s_(*t*) and *μ*_d_(*t*) in the system (25)–(36). The parameters *σ*_ss_,*σ*_ds_,*σ*_sd_ and *σ*_dd_ remain constant.

For mathematical convenience we write the modulations in complex form

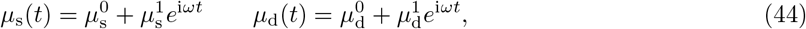

with small 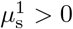 and 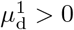 (thereby introducing a companion system) and approximate the solution to first order, 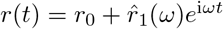, where 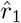 is a complex variable from which the response amplitude *r*_1_ and phase shift *ψ* of the oscillatory spike rate *r*_0_ + *r*_1_(*ω*) sin(*ωt* + *ψ*(*ω*)) can be extracted in a straightforward way: 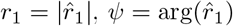 (see, e.g., [45] for a similar type of analysis in a different setting). Note that also the state variables of this solution take the form 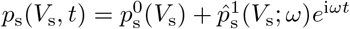 (analogously for *p*_s,1_, *p*_s,2_, *u*_s_, *u*_s,1_, *u*_s,2_). For fixed (angular) frequency ω we obtain the following ODE system (neglecting terms of second and higher order in 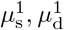):

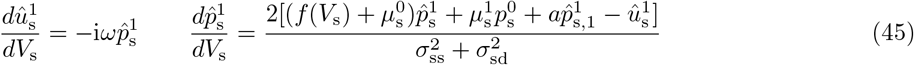

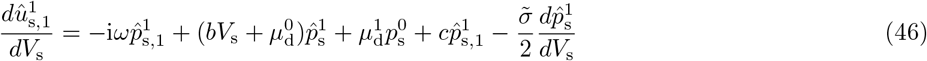

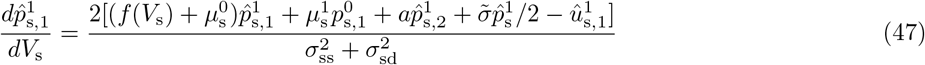

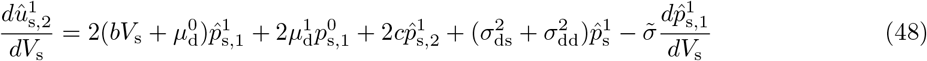

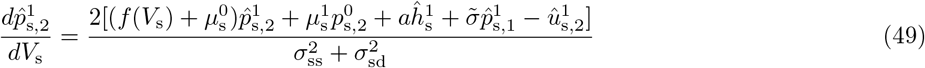

with

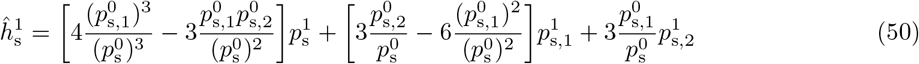

subject to the conditions

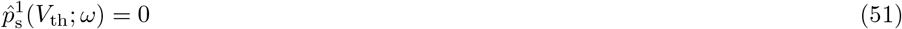

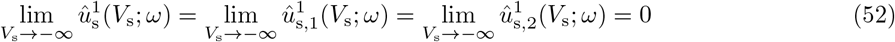

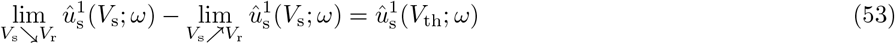

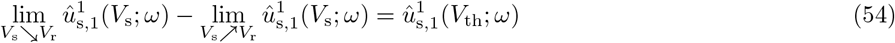

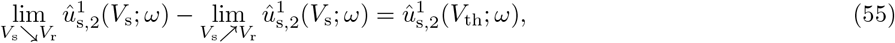

where 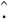 indicates a complex valued variable that depends on ω. Note that this ODE system depends on the (steady state) solution of the system (39)–(43) through 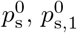 and 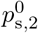.

*Numerical solution* The linear (complex valued) ODE system with variable coefficients (45)–(55) can be conveniently solved in the following way. The desired spike rate response solution can be written as 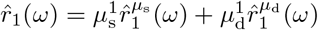 where the solution component 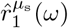 corresponds to 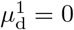 and, vice versa, 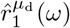 corresponds to 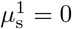. We first describe how we obtain 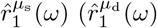 is calculated in an analogous way, see further below). The solution 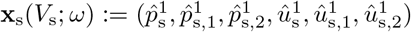 associated with 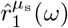 can be decomposed into (omitting the argument *ω* below for improved readability)

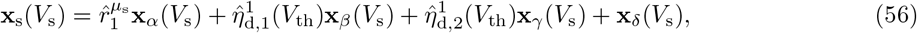

where x_*α*_, x_*β*_, x_*γ*_ solve the homogeneous part 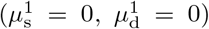 of the ODE system (45)–(50) with 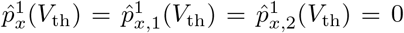 for *x* ∈ {*α,β,γ*} 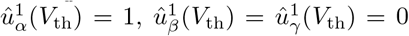, 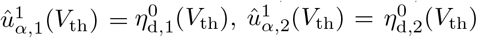, 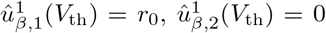, 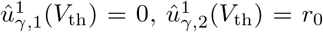, and “jump” conditions (53)–(55). Note that 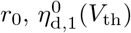 and 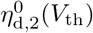 are known from the solution for the steady state system. x_*δ*_ solves the inhomogeneous system (45)–(50) with 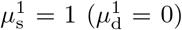 and condition x_*δ*_(*V*_th_) = 0. These solutions are obtained numerically by backward integration from *V*_th_ to *V*_lb_. 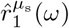 together with 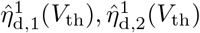 are then calculated by solving the linear equation system that arises to satisfy the condition 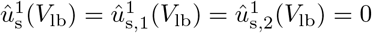. The solution method for 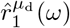 is completely analogous with the difference that 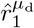 appears instead of 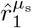 in Eq. (56) and x_*δ*_ solves the inhomogeneous system (45)–(50) with 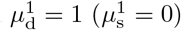.

We obtain the amplitude of the spike rate response to an applied field from

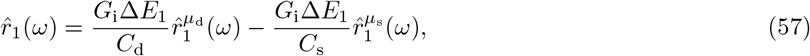

whereas the response modulations to sinusoidal mean input at the soma and dendrite in the absence of an oscillatory field are given by 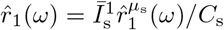 and 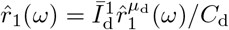, respectively.

### 4. Parametrization via ball-and-stick model

In the following we describe a semi-analytical technique to fit the two-compartment (2C) model to a biophysically more detailed, spatially extended ball-and-stick (BS) model. In particular, the parameter values of the 2C model are determined to best approximate the somatic voltage dynamics of the BS model. This is done in an efficient way using analytical results for the voltage dynamics of both models, and it does not depend on a particular choice of parameter values for the input or the extracellular field. This part may thus be regarded as a reduction of the BS model.

#### Ball-and-stick model

The BS neuron model consists of a finite passive dendritic cable of length *L* with lumped somatic compartment at the proximal end, *x* = 0, and a sealed end boundary condition at the distal extremity, *x* = *L*. It includes capacitive and leak currents across the membrane, an approximation of the spike-generating sodium current at the soma, an internal current (along the cable) and synaptic input currents at the soma and distal dendrite as well as a spatially homogeneous but time-varying external electric field (for details on the derivation of this model see [20]). The dynamics of the model are governed by

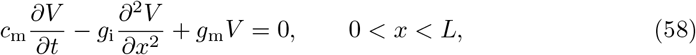

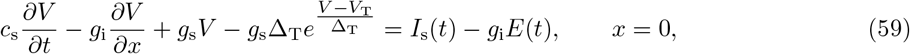

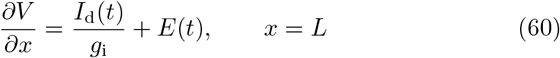

together with a reset condition

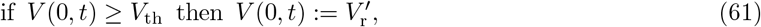

where *c*_m_ = *cD*_d_*π* is the membrane capacitance, *g*_m_ = *ϱ*_m_*D*_d_*π* is the membrane conductance and *g*_i_ = *ϱ*_i_(*D*_d_/2)^2^*π* is the internal (axial) conductance of a dendritic cable segment of unit length. *c* is the specific membrane capacitance (in F/m^2^), *ϱ*_i_ is the specific internal conductance (in S/m), *ϱ*_m_ is the specific membrane conductance (in S/m^2^) and *D*_d_ is the cable diameter. 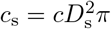 and 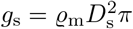 are the somatic membrane capacitance and leak conductance, respectively, with soma diameter *D*_s_. The exponential term with threshold slope factor Δ_T_ and effective threshold voltage *V*_T_ approximates the rapidly increasing Na^+^ current at spike initiation [40]. Spike times are defined by the times at which the somatic membrane voltage *V*(0,*t*) crosses the threshold voltage value *V*_th_ from below (cf. spike mechanism of the 2C model). *I*_s_(*t*), *I*_d_(*t*) and *E*(*t*) are described by Eqs. (8), (9) and (7), respectively. The parameter values are provided in table 1.

**Table 1.** Description and values of the ball-and-stick model parameters

| Parameter (unit) | Description | Value |
| --- | --- | --- |
| $c$ ($\text{F/m}^2$) | Specific membrane capacitance | $1 \cdot 10^{-2}$ [36, 46] |
| $\varrho_m$ ($\text{S/m}^2$) | Specific membrane conductance | $1/3$ [46] |
| $\varrho_i$ ($\text{S/m}$) | Specific internal conductance | $1/2$ [47] |
| $D_s$ (m) | Soma diameter | $15 \cdot 10^{-6}$ [48] |
| $D_d$ (m) | Dendritic cable diameter | $1 \cdot 10^{-6}$ [47] |
| $L$ (m) | Dendritic cable length | $7 \cdot 10^{-4}$ [49] |
| $\Delta_T$ (mV) | Threshold slope factor | $1.5$ [50] |
| $V_T$ (mV) | Effective threshold voltage | $10$ [50] |
| $V_{\text{th}}$ (mV) | Threshold (spike) voltage | $20$ |
| $V_r'$ (mV) | Reset voltage | $0$ |

To generate spike trains we simulated the BS neuron model using a semi-implicit numerical scheme (Crank-Nicolson method; see, e.g., appendix C of [51]) that was extended for stochasticity as proposed in [52], and by applying the tridiagonal matrix algorithm. Discretization steps were 5 *μ*s for time and *L*/200 for space (along the dendrite).

#### Calculation of subthreshold responses

We analytically calculate the somatic membrane voltage response of the BS model for small variations of the synaptic inputs *I*_s_(*t*), *I*_d_(*t*) and a weak oscillatory field *E*(*t*), which do not elicit spikes. We consider that the somatic voltage evolves sufficiently below *V*_T_, which allows us to neglect the exponential term in Eq. (59) (cf. Methods section 2). The linear PDE (58) together with the boundary conditions (59) and (60) is then solved using separation of variables *V*(*x,t*) = *W*(*x*)*U*(*t*) and the Fourier transform

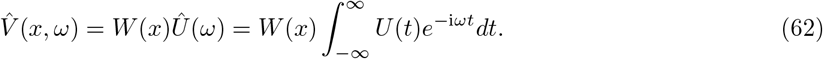

We obtain the system of differential equations

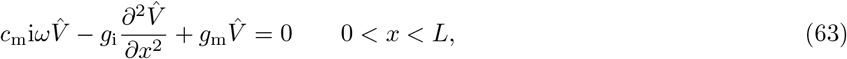

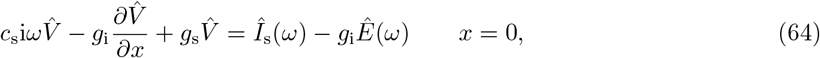

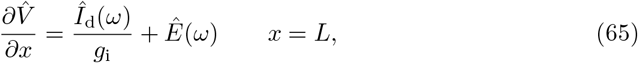

where 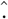 indicates a (temporally) Fourier transformed variable. The solution of this system can be expressed as

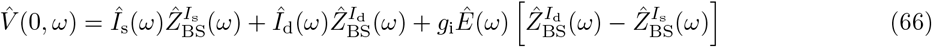

with

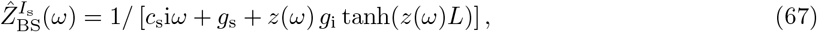

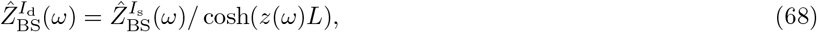

where ±*z*(*ω*) are the roots of the characteristic polynomial *g*_i_*y*^2^ = *g*_m_ + *c*_m_i*ω* of Eq. (63),

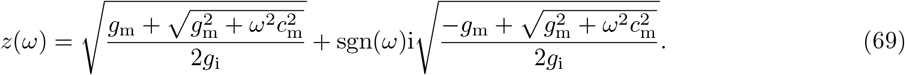

Note that 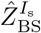 and 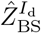 are the somatic impedances for inputs at the soma and the distal dendrite, respectively.

The response to a sinusoidal field variation, *E*(*t*) = *E*_1_ sin(*φt*), with constant *I*_s_ and *I*_d_ can be expressed in the time domain as

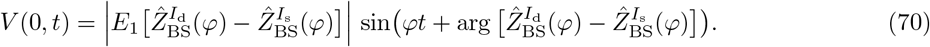

In addition we calculate the somatic voltage time series in response to subthreshold input for a given initial condition *V*(*x*, 0) = *V*_0_(*x*). Using separation of variables and the Laplace transform

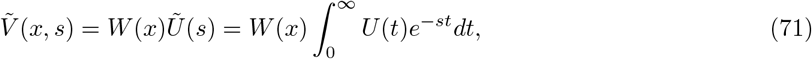

with complex variable s we obtain

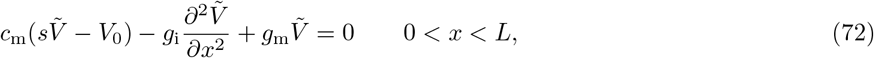

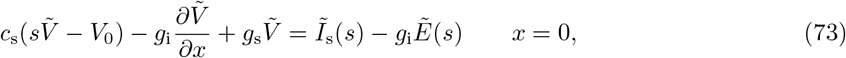

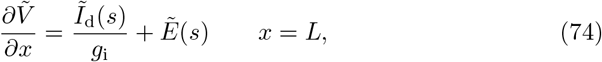

where 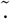 indicates a (temporally) Laplace transformed variable. We solve this system and obtain

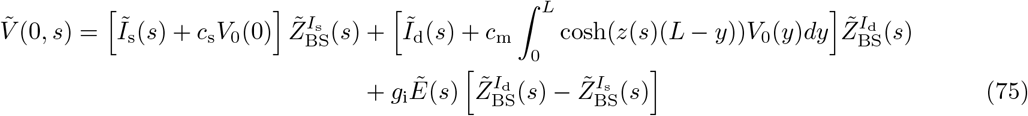

with

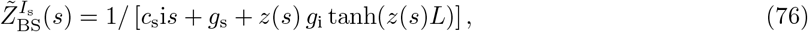

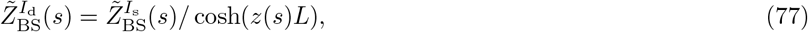

where ±*z*(*s*) are the roots of the characteristic polynomial *g*_i_*y*^2^ = *c*_m_*s* + *g*_m_ of Eq. (72), given by

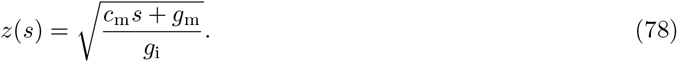

The somatic voltage time series *V*(0, *t*) is then computed by inverse transforming *Ṽ*(0, *s*) using an efficient numerical method [53].

#### Parameter fitting

We approximate the somatic voltage dynamics of the BS model by the 2C model in two steps. First, we fit 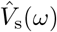 to 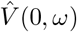 using Eqs. (10) and (66) over a range of angular frequencies *ω* ∈ [0, *ω*_max_], requiring that the voltage values for *ω* = 0 (i.e. the steady states) match exactly. This constraint determines three parameters,

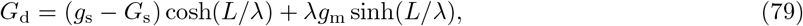

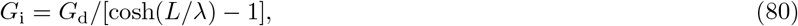

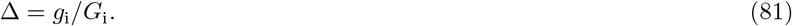

where we have introduced the electrotonic length constant 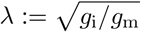. The remaining three subthreshold parameters *C*_s_, *C*_d_ and *G*_s_ are obtained using the method of least squares, with *ω*_max_/(2*π*) = 10 kHz. In the second step we determine the reset voltage *V*_r_ by approximating the transient BS somatic voltage time series immediately after a spike elicited by threshold somatic input and threshold dendritic input for *E*(*t*) = 0. Specifically, we fit the post-spike voltage time series *V*_s_(*t*) to *V*(0, *t*) across the time interval *t* ∈ [0, *τ*_m_] with initial conditions *V*_s_(0) = *V*_r_, *V*_d_(0) = (*G*_i_*V*_th_ + *I*_d_)/(*G*_d_ + *G*_i_) and 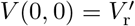,

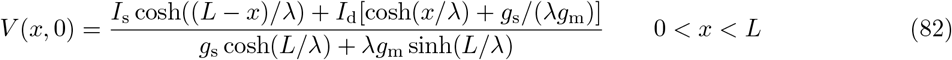

for threshold somatic input *I*_s_ = *V*_th_[*g*_s_ + *λg*_m_ tanh(*L*/*λ*)], *I*_d_ = 0 and for threshold dendritic input *I*_d_ = *V*_th_[*g*_s_ cosh(*L*/*λ*)+*λg*_m_ sinh(*L*/*λ*)], *I*_s_ = 0 simultaneously using the method of least squares. *τ*_m_ = *C*_s_/(*G*_s_+*G*_i_) is the somatic membrane time constant of the 2C model. Note that we consider threshold input as constant current that yields *V* = *V*_T_ (calculated from the linear subthreshold model systems). The voltage time series *V*(0, *t*) of the BS model is rapidly computed using the Laplace transform, Eq. (75), and the voltage time series *V*_s_(*t*) of the 2C model is calculated analytically in a straightforward way (linear ODE system). To guarantee an equal effectiveness of the exponential term on the somatic membrane voltage dynamics in the 2C model compared to the BS model we set *G_e_* = *C*_s_*g*_s_/*c*_s_ (cf. Eqs. (1) and (59)). The values for Δ_T_, *V*_T_ and *V*_th_ are set equal to those of the BS model. Notably, this fitting method is very efficient, since it involves analytical results and does not depend on specific realizations of time series for the neuronal input or extracellular field.

### 5. Spike coincidence measure

To quantify the similarity between the spike trains of the two-compartment and the ball-and-stick model neurons we used the coincidence factor Γ defined by [54]

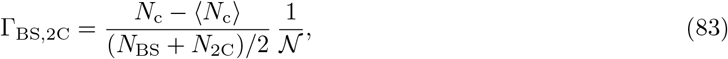

where *N*_c_ is the number of coincident spikes with precision (i.e., maximal temporal separation) Δ_c_, *N*_BS_ and *N*_2C_ are the number of spikes in the spike trains of the ball-and-stick and the two-compartment models, respectively. 〈*N*_c_〉 = 2*r*Δ_c_*N*_BS_ is the expected number of coincidences generated by a homogeneous Poisson process with spike rate *r* = *N*_2C_/*T* as exhibited by the two-compartment model, where *T* is the duration of the spike train. The factor 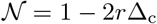 normalizes Γ_BS,2C_ to a maximum value of 1, which is attained if the spike trains match optimally (with precision Δ_c_). Γ_BS,2C_ = 0 on the other hand would result from a homogeneous Poisson process with rate that corresponds to the spike train of the two-compartment model, and therefore indicates pure chance. Here we used Δ_c_ = 5 ms.

### 6. Two-population mean-field network

In this section we derive a mean-field network model from a large number *N* of sparsely and randomly coupled pyramidal (PY) neurons and inhibitory (IN) interneurons. Each PY neuron is described by the 2C model, Eqs. (1)–(4), (7)–(9), and each IN neuron is described by an exponential integrate-and-fire (point) neuron model,

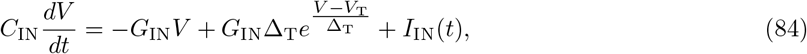

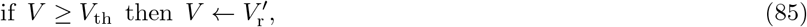

because IN neurons do not exhibit an elongated spatial morphology compared with PY neurons. We used *C*_IN_ = 0.2 nF and *G*_IN_ = 10 nS. The model neurons receive fluctuating external and recurrent synaptic input and are exposed to an applied weak electric field *E*(*t*) that is spatially uniform. For fields induced by transcranial brain stimulation [7] this is a valid assumption. Each PY neuron receives inputs from *K*_s_ IN neurons at the soma and *K*_d_ IN neurons at the dendrite and each IN neuron receives inputs from K_IN_ PY neurons. Synaptic coupling is described by delayed current pulses that produce postsynaptic potentials of size J_s_, *J*_d_ or *J*_IN_ (depending on the location of the synapse). Specifically, the input currents for neuron *k*, with somatic and dendritic membrane voltage *V*_s,*k*_, *V*_d,*k*_ (in case of a PY neuron) or overall membrane voltage *V_k_* (in case of an IN neuron), are given by

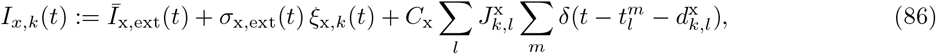

x ∈ {s, d, IN}, where 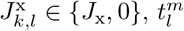 is the *m*-th spike time of neuron *l* and 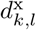 the coupling delay between neuron *l* and *k*. The delays are independently sampled according to the probability distribution with density

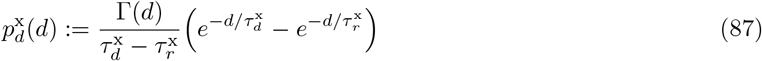

x ∈ {s, d, IN}. We consider large numbers of connections *K*_x_ and reasonably small coupling strengths *J*_x_.

For large networks (in the mean-field limit *N* → ∞) the overall synaptic input can be approximated by a mean part with additive fluctuations,

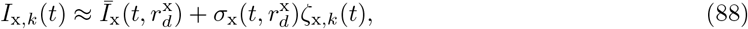

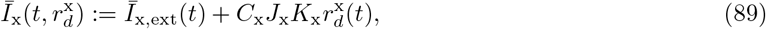

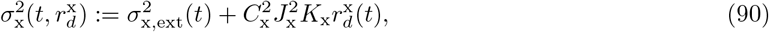

x ∈ {s, d, IN} with delayed spike rates 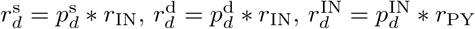, and unit white Gaussian noise process *ζ*_x,*k*_ that is uncorrelated to that of any other neuron (see, e.g., [27]). This step is valid under the assumptions of (i) sufficient presynaptic activity, (ii) that neuronal spike trains can be approximated by Poisson processes and (iii) that the correlations between the fluctuations of synaptic inputs for different neurons vanish. Note that the latter assumption is supported by sparse and random synaptic connectivity.

#### Fokker-Planck based description and resonance analysis

The previous approximation allows us to express the collective spiking dynamics in terms of a coupled system of Fokker-Planck PDEs, one for the PY population (cf. Methods section 3) and one for the IN population, where the coupling is mediated through the synaptic input moments *Ī*_x_ and 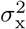 (cf. Eqs. (89) and (90); see, e.g., [27]). To analyze the resonance properties of this mean-field network model we consider either a weak sinusoidal electric field, Eq. (7), or weak sinusoidal modulations of the external mean input at the soma or dendrite of PY neurons, 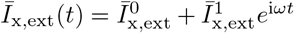, x ∈ {s, d}. We assume a parametrization of the network such that without these modulations 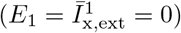 it exhibits asynchronous activity represented by a fixed point solution of the mean-field system.

We write the modulations in complex form and express the first order population spike rate responses as (cf. Methods section 3) 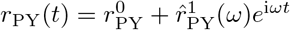 and 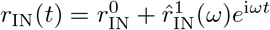. To obtain the stationary (steady state) components 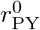 and 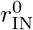 we solve the system (39)–(43) with 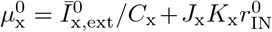 and 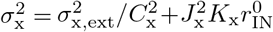, x ∈ {s, d} (and with the corresponding boundary conditions) together with the respective system for the IN population with 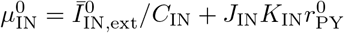 and 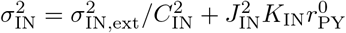 via fixed point iteration of the form 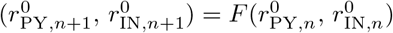. Note that *σ*_sd_ = *σ*_ds_ = 0 in Eqs. (13) and (14). The response components 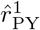 and 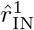 are obtained from

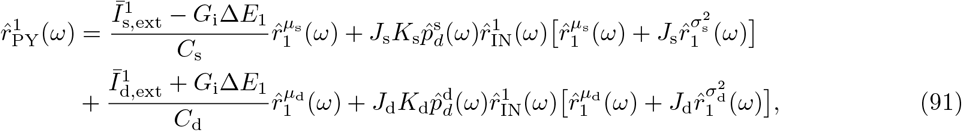

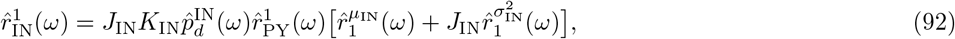

where 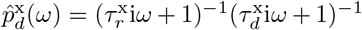, x ∈ {s, d, IN} is the Fourier transformed delay density (cf. Eq. (87)) and 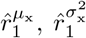 are the population spike rate response components for sinusoidal modulations of *μ*_x_, 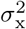 of unit amplitude. Note that Eqs. (91) and (92) can be jointly solved in a straightforward way. In addition to first order responses to modulations of *μ*_s_ and *μ*_d_ we therefore need to calculate the responses for weak sinusoidal modulations of 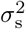 and 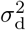. That is, the rate response solution of the system (25)–(36) for 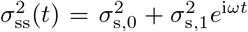 and 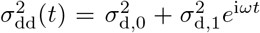 is required. This is done in an analogous way as explained above for sinusoidal modulations of *μ*_s_ and *μ*_d_ (see Methods section 3): we solve (45)–(55) with 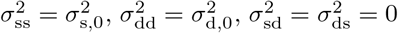, and where the inhomogeneous terms 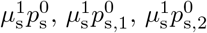 and 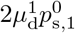 in Eqs. (45), (47), (49) and (48) are replaced by 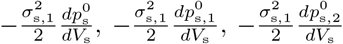 and 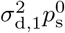, respectively. The resulting system can be numerically solved as explained in Methods section 3, using Eq. (56) where 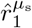 is replaced by 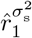 and 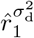, respectively, and x_*δ*_ solves the adjusted inhomogeneous system (with 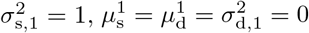 and 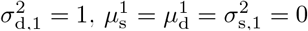, respectively). For the simpler case of the exponential integrate-and-fire point model for the IN population (steady state and response modulations) see [27, 45].

## Acknowledgments

This work was supported by Deutsche Forschungsgemeinschaft in the framework of Collaborative Research Center 910. The funders had no role in study design, analysis, decision to publish, or preparation of the manuscript.

## Author Contributions

J.L. conceived the study, developed the analytical methods and wrote the initial draft of the manuscript. J.L. and K.O. reviewed and edited the manuscript, and acquired funding for the project.

